# A Novel Allosteric Inhibitor Targeting IMPDH2 at Y233 Overcomes Resistance to Tyrosine Kinase Inhibitors in Lymphoma

**DOI:** 10.1101/2025.08.19.667179

**Authors:** Nagarajan Pattabiraman, Cosimo Lobello, David Rushmore, Luca Mologni, Mariusz Wasik, Johnvesly Basappa

**Affiliations:** Veracure Biosciences Inc., Silver Spring, MD; Department of Pathology, Fox Chase Cancer Center, Philadelphia, PA, USA; Department of Medicine and Surgery, University of Milano-Bicocca, Monza, Italy

**Keywords:** *IMPDH2*, PI3P, tyrosine phosphorylation, *In-silico* screening, IMPDH drug discovery

## Abstract

Inosine monophosphate dehydrogenase-2 (IMPDH2) is a rate-limiting enzyme in the de novo biosynthesis of guanine nucleotides and is often overexpressed in hematologic malignancies and solid tumors. However, its regulatory mechanisms in cancer remain poorly understood. Here, we identify IMPDH2 as a direct substrate of the oncogenic kinases ALK and SRC, which phosphorylate tyrosine 233 (Y233) within the enzyme’s allosteric Bateman domain. Using peptide-based mutagenesis and enzymatic assays, we demonstrate that Y233 phosphorylation is essential for IMPDH2 activity. We found that phosphoinositide-3-phosphate (PI3P), a signaling phospholipid, selectively binds to and inhibits IMPDH2, but not its isoform IMPDH1, revealing a novel lipid-mediated, isoform-specific regulatory mechanism. Next, we conducted structure-based virtual screening and identified a potent allosteric inhibitor of IMPDH2, compound 10 (comp-10), with an IC₅₀ of 260 nM. Comp-10 significantly impairs cell proliferation in ALK-positive anaplastic large cell lymphoma (ALCL) cell lines, including those resistant to the ALK inhibitors crizotinib and lorlatinib, and outperformed the FDA-approved IMPDH inhibitor mycophenolic acid. These findings reveal the dual regulation of IMPDH2 through tyrosine phosphorylation and binding to PI3P, and describe the discovery of a new IMPDH2 inhibitor, suggesting a potential therapeutic strategy to overcome resistance to tyrosine kinase inhibitors.

## Introduction

Humans and other mammals express two IMPDH isoforms, IMPDH1 and IMPDH2, each comprising 514 amino acids and sharing 84% sequence identity. Both isoforms contain a catalytic domain that binds substrates and a regulatory Bateman domain, which modulates enzymatic activity through allosteric interactions with the catalytic core(1). The IMPDH regulatory (Bateman) domain contains ATP- and GTP-binding sites. IMPDH exists as a constitutive tetramer, and nucleotide binding promotes reversible dimerization of the regulatory domains, leading to octamer formation(2). IMPDH1 is constitutively expressed in normal lymphocytes but is highly upregulated in a subset of small cell lung cancers (SCLC)(3). IMPDH2 is overexpressed in hematological malignancies, including human leukemic cell lines and BCR-ABL–positive acute myelogenous leukemia(4), in chronic myelogenous leukemia(*5*) and other cancers, such as triple-negative breast cancer(6), prostate cancer(7, 8), kidney cancer(9), nasopharyngeal carcinoma(10), in a subset of small-cell lung cancers(3, 11), in non-small cell lung cancer(12) and in glioblastoma(13, 14) and brain metastases(15). Proteomic profiling of colorectal cancer plasma identified IMPDH2 as a potential biomarker(16). Whereas protein tyrosine phosphorylation makes up only 2.5%, it has significant effects on nearly every aspect of cellular physiology(17). Our previous work demonstrated that ALK and SRC kinase-mediated tyrosine phosphorylation of ACLY regulates its function(18, 19). Although IMPDH2 overexpression in solid tumors is well documented, its post-translational regulation, particularly by tyrosine phosphorylation remains poorly understood. Here, we report for the first time that IMPDH2 is phosphorylated on a critical tyrosine residue by oncogenic kinases, as demonstrated through in vitro kinase assays and mass spectrometry-based phosphoproteomic analysis.

IMPDH1 and IMPDH2 pose a major challenge for isoform-specific drug development due to their 84% sequence identity. To overcome this, we analyzed sequence differences and hypothesized that the two isoforms may possess distinct phosphoinositide (PI) binding sites. PIs are lipid second messengers critical for membrane trafficking, metabolism, growth, signaling, and autophagy. Their phosphorylation generates seven distinct species, including PI3P, a key marker of endosomal and autophagic membranes. PI3P is recognized by FYVE and PX domain-containing proteins, suggesting that isoform-specific PI interactions could be leveraged for selective targeting of IMPDH isoforms(20, 21). FDA-approved IMPDH inhibitors— such as Mycophenolic acid (MPA), Mycophenolate mofetil (MMF), Ribavirin, and Mizoribine target the catalytic domains of both IMPDH1 and IMPDH2 and are used clinically for immunosuppressive and antiviral therapy(22). These inhibitors also promote the formation of Rods and Rings (RRs), or IMPDH filaments, which are non-membrane-bound intracellular polymeric structures(23–25).

Our study shows that mycophenolic acid (MPA) induces expression of catalytically inactive IMPDH2 and promotes filament formation in T-cell and B-cell lymphomas, as well as in solid tumors. This filamentous assembly may contribute to the toxicity and off-target effects of current IMPDH inhibitors, highlighting the need for more selective therapeutics. We identify IMPDH2 as a direct substrate of the oncogenic kinases ALK and SRC, and demonstrate that PI3P binding selectively inhibits IMPDH2 activity but not that of IMPDH1. A self-derived IMPDH1 peptide from the Y233 domain also suppresses IMPDH2 function. Based on these findings and structural modeling, we performed in silico screening and discovered a novel allosteric inhibitor, comp-10, which specifically targets the allosteric domain and is distinct from current IMPDH inhibitors, thereby opening avenues for future therapeutic advancements.

## Materials and methods

### Reagents and antibodies

Recombinant human IMPDH1 (Catalog # 8904-DH) and IMPDH2 (catalog# Catalog # 8349-DH) proteins were obtained from R&D Systems. IMPDH activity assay kit was from Biovision (Catalog#K495) or abcam (catalog#ab283395). PIP Strips – Lipid-Protein Interaction Assay (catalog#P-6001), PIP Arrays – Lipid-Protein Interaction Assay (catalog# P-6001), PI(3)P Beads (catalog#P-B003A), PI(4,5)P2 diC4 (catalog#P-4504) and PI(3)P diC8 (catalog#P-3008A) were purchased from Echelon Biosciences. All other reagents are from Sigma-Aldrich. Cell Proliferation Reagent WST-1 (Millepore-sigma), CellTiter 96® Non-Radioactive Cell Proliferation Assay (MTT) kit (catalog#G4001) from Promega. HA Tag Monoclonal Antibody ( Thermo Fisher Scientific Cat#26183, RRID: AB_10978021), Pierce IP Lysis Buffer (Thermo Fisher Scientific Cat#87787), Pierce Anti-HA Magnetic Beads ( Thermo Fisher Scientific Cat# 88837, RRID: AB_2861399) Halt Protease and Phosphatase Inhibitor (Thermo Fisher Scientific Cat#78440), ALK Recombinant Human Protein (Thermo Fisher Scientific, Cat# PV3867) and SRC Recombinant Human Protein (Thermo Fisher Scientific, Cat# P3044). Mizoribine (Cat #S1384), Ribavarin (Cat #S2504), mycophenolic acid (Cat# S2487), Mycophenolate mofetil (Cat#S1501), certinib (Cat#S7083). All these small-molecule inhibitors were purchased from Selleckchem. IMPDH1 (RRID:AB_2878992, Cat# 22092-1-AP), IMPDH2 (RRID:AB_2127351, Cat# 12948-1-AP) and GAPDH (RRID:AB_2107436, Cat# 60004-1-Ig) from Proteintech, HA-Tag (C29F4) Rabbit (RRID:AB_1549585, Cat# 3724S), Phospho-ALK (Tyr1604) Antibody (RRID:AB_331047, Cat# 3341S), ALK (D5F3^®^) XP^®^ Rabbit (RRID:AB_11127207, Cat# 3633S) from Cell Signaling Technology.

### IMPDH1 and IMPDH2 peptide synthesis

Peptide-I (Human IMPDH1): (aa122-136), SPSHTVGDVLEAKMR; Peptide-I: (Human IMPDH1-WT): (aa228-242), KKNRD**Y**PLASKDSQK; Peptide-I: (Human IMPDH1-Y233F): (aa228-242), KKNRD**F**PLASKDSQK; Peptide-II (Human IMPDH2): (aa122-136), SPKDRVRDVFEAKAR; Peptide-II (Human IMPDH2-WT): (aa228-242), KKNRD**Y**PLASKDAKK and Peptide-II (Human IMPDH2-Y233F): (aa228-242), KKNRD**F**PLASKDAKK were synthesized at GenScript, Piscataway, NJ.

### Plasmids and lentivirus products

Custom Plasmid Preparation: IMPDH2-HA tagged_pLenti_MS2-P65-HSF1_mCherry, Custom Plasmid Preparation: IMPDH2-HA tagged_pLenti_MS2-P65-HSF1_GFP and Express Mutagenesis: IMPDH2-HA tagged Y233F_pLenti_MS2-P65-HSF1_GFP synthesized at GenScript, Piscataway, NJ. pLenti-EGFP-2xFYVE (Plasmid #136996, pLenti-EGFP-2xFYVE was a gift from Ken-Ichi Takemaru (Addgene plasmid Cat#136996; http://n2t.net/addgene:136996;RRID:Addgene_136996), 3rd Gen. Packaging Mix & Lentifectin Combo Pack from Applied Biological System (ABM good# LV053-G074), Lenti-X™ Concentrator from TAKARA (catalog# 631231).

### MCL, DLBCL, and ALCL cell lines

MCL, MCL-RL cells were derived from a patient with MCL at the University of Pennsylvania, Philadelphia, PA. JeKo-1, Maver Rec-1, Granta519, DLBCL, OCI-LY1, OCI-LY4, OCI-LY8, and TOLEDO; SUDHL-1, JB6, Karpas 299, SUP-M2, L82, and SR786 cell lines were derived from ALK+ALCL patients and cultured as described in earlier(26). Human CD4+ cells transduced with NPM-ALK, NA1, were created by our group as described earlier(27). The cell lines were regularly tested for Mycoplasma contamination using Mycoplasma detection kits from Thermo Fisher Scientific. All cell lines were routinely tested for Mycoplasma contamination using Mycoplasma detection kits (Thermo Fisher Scientific) and were authenticated. Cells were maintained in RPMI-1640 medium supplemented with 10% fetal bovine serum (FBS) and 1% penicillin-streptomycin (Pen/Strep) in a humidified incubator at 37°C with 5% CO₂.

### ALK inhibitor (ALKi) resistant ALCL cell lines

ALKi crizotinib-resistant cell lines described earlier, KARPAS299 CR06 (resistant up to 600 nM of crizotinib and cultured at that concentration) and SUPM2 CR03 (resistant up to 300 nM of crizotinib and cultured at that concentration), and ALKi lorlatinib-resistant cell lines described earlier, KARPAS LR1000 (resistant up to 1000 nM of lorlatinib and cultured at 300 nM of lorlatinib) and SUPM2 LR1000 (resistant up to 1000 nM of lorlatinib and cultured at 300 nM of lorlatinib). All of these cell lines were obtained from Drs. Carlo Gambacorti-Passerini and Luca Mologni, University of Milano-Bicocca, Italy(28). The cell lines were grown in RPMI medium supplemented with 10% FBS and 1% penicillin/streptomycin in a humidified incubator at 37°C with 5% CO2.

### IMPDH1 and IMPDH2 PIP strip and PIP array binding assay

For initial experiments, human recombinant IMPDH1 and IMPDH2 proteins were diluted to 1 µg/ml in 3% BSA and dissolved in PBS buffer. Next, the IMPDH2 concentration was further diluted to 100 ng/ml. The diluted proteins were subjected to a PIP strip lipid-protein interaction assay as described earlier. Based on IMPDH2’s unique binding to PI3P compared to IMPDH1, we further validated IMPDH2’s binding to PI3P on a PIP array coated with various concentrations.

### IMPDH1 and IMPDH2 activity assay in the presence of PIP2 (PI(4,5)P2) and PI3P (PI(3)P diC8)

Activity assays on human recombinant IMPDH1 and IMPDH2 were conducted using a Synergy H1 microplate reader from BioTek, 96-well plate readers, and a commercially available kit from Biovision, Inc. According to the assay kit instructions, the phospholipids PIP2 and PI3P were diluted in activity assay buffer and preincubated with recombinant IMPDH1 or IMPDH2 at concentrations ranging from 50 µM to 200 µM for 15 minutes. After this incubation, a substrate solution was added to the wells to initiate enzyme activity. The activity was continuously measured every minute until 30 to 60 minutes later triplicate.

### IMPDH1 and IMPDH2 Enzyme inhibition assays in the presence of peptides

Activity assays on human recombinant IMPDH1 and IMPDH2 were conducted using SynergyH1 microplate reader BioTek,96-well plate readers, and a commercially available kit from Biovision, Inc. As outlined in the assay kit instructions, the peptides synthesized by GenScript were first incubated with recombinant IMPDH1 or IMPDH2 enzymes for 30 minutes. Following this incubation, a substrate solution was added to the wells to initiate enzyme activity, with each peptide and control being tested in triplicate.

### *In-vitro* kinase assay and LC-MS/MS phosphoproteomics analysis

We performed an *in-vitro* kinase assay on human recombinant IMPDH2 in the presence of active anaplastic lymphoma kinase (ALK) and SRC kinase. We subjected it to LC-MS/MS analysis as we described in our previous study(18).

### Structure-based *In-silico* screening of IMPDH inhibitors

The National Cancer Institute (NCI) maintains a database of compounds, known as the Mechanistic Set VI, which comprises 811 compounds derived from the 37,836 open compounds tested in the NCI human tumor 60-cell line screen. This mechanistic diversity set was chosen to represent a broad range of growth inhibition patterns in the NCI60 cell line screen based on the GI50 activity of the compounds. Compounds tested in the NCI-60 cell line screen were clustered using the FASTCLUS procedure in the SAS statistical package. This algorithm is based on MacQueen’s k-means algorithm, which minimizes the sum of squared distances from the cluster means. The procedure resulted in 1272 clusters. A single representative compound from each cluster, for which an adequate supply of material was available, was chosen. Some clusters are not represented in the set, as insufficient material was available. The database of compounds was downloaded. We subjected the compounds in the database with the following filter: 1) 100 <= Molecular Weight =< 500 2) Only one molecule, 3) 2 <= Number of rotatable bonds =< 6 4) Lipenski druglike =1 and no chiral centers. This resulted in 283 compounds for in silico screening. We generated ∼15,200 energetically favorable conformations for *in-silico* screening. We docked these conformations into the GTP binding site of IMPDH2 and ranked them in order based on a scoring function. We selected 100 top-scoring compound-IMPDH2 complexes and selected 38 as the unique compounds based on National Service Center (NSC) identity. We got 15 compounds from the National Cancer Institute (NCI) and tested them for enzyme activity inhibition against recombinant human IMPDH2 in 96-well plate screening.

### IMPDH2 self-derived peptide docking to human IMPDH2

We analyzed the structural details of 17 Electron microscopic, 5 X-ray, and one AlphaFold structure of IMPDH2 from PDB (https://www.rcsb.org), and we chose the Alphafold structure (AF-P12268-F1) for in-silico docking studies of 15mer peptide derived from IMPDH2. The sequence of the 15mer peptide from IMPDH2 is 228-**KKNRDYPLASKDAKK**-242. Since the 15mer contains proline at position 234, which restricts the conformation of neighboring residues, and in addition to the potential salt bridges between acidic and basic amino acids, the 3D structure of the 15mer peptide is not very flexible. The coordinates of the 3D structure of the 15mer peptide were taken from the 228-242 region of the AlphaFold structure. The peptide structure was subjected to energy minimization using the AMBER-EHT force field.

### Cell proliferation assay

For the cell proliferation assay, the aforementioned cell lines were plated in 96-well plates at a density of 20,000 to 40,000 cells per well in standard RPMI medium, with DMSO as a control or with the respective drug concentrations in 100 µL of medium. After 48 hours in culture, cell proliferation was assessed using the WST-1 or MTT assay method following the manufacturer’s protocol.

### Colony formation assay

The colony formation assay used Human Methylcellulose Complete Media (Catalog #: HSC003, R&D Systems). In brief, 200 cells per well in triplicate (in a 6-well plate) were placed in Human Methylcellulose media according to the manufacturer’s protocol, with a DMSO control, comp-10 (100 nM), and MPA (100 nM). After plating the cells, the plate was incubated in a standard cell culture chamber for four weeks. Colony formation was then visualized using the iBright imaging system, 1500 (Thermo-Fisher Inc.), and the number of cells was counted.

### Statistics and reproducibility

The student’s *t*-test was used to analyze differences in Western blot densitometric values and activity assays to assess differences in cell growth and colony formation. *p-*values equal to or less than 0.05 were considered statistically significant without being adjusted for multiple comparisons. The statistical analysis was performed using GraphPad Prism 9.0. software and NIH ImageJ software for the densitometric quantitation of western blot data.

## Data availability

We confirm that all relevant data and methods are included in the main Article and the Supplementary Information section.

## Results

### IMPDH2 is significantly overexpressed in hematological malignancies

IMPDH1 and IMPDH2 are rate-limiting enzymes critical for purine nucleotide biosynthesis, particularly during periods of rapid cell proliferation, such as in activated T and B cells. While IMPDH1 is constitutively expressed across most tissues, IMPDH2 is highly inducible and is frequently overexpressed in cancer. The two isoforms share approximately 85% sequence similarity (Supplementary Fig. 1). Analysis of publicly available RNA-seq data from the Human Protein Atlas revealed widespread overexpression of IMPDH2 across a broad range of cancer cell lines compared to IMPDH1(Supplementary Fig. 2a, b). In hematological malignancies including mantle cell lymphoma (MCL), diffuse large B-cell lymphoma (DLBCL), chronic lymphocytic leukemia (CLL), acute myeloid leukemia (AML), and anaplastic large cell lymphoma (ALCL) IMPDH2 expression was markedly higher than IMPDH1 (Supplementary Fig. 2c). Similarly, in leukemia and multiple myeloma cell lines, IMPDH2 remained the dominant isoform with minimal expression of IMPDH1 (Supplementary Fig. 2e–h). These findings underscore the cancer-associated upregulation of IMPDH2 and support its prioritization as a therapeutic target.

### *In-vitro* kinase and phosphoproteomic analyses reveal tyrosine phosphorylation of IMPDH2 in the allosteric domain

IMPDH1 and IMPDH2 are highly expressed in proliferative lymphoid malignancies, including T- and B-cell lymphomas. In this study, we focused on two aggressive subtypes—anaplastic lymphoma kinase (ALK)-positive anaplastic large cell lymphoma (ALCL) and mantle cell lymphoma (MCL). Given the known role of tyrosine phosphorylation in modulating metabolic enzymes in tyrosine kinase–driven cancers, we explored whether oncogenic tyrosine kinases phosphorylate IMPDH2. Although public databases such as the Human Protein Atlas and PhosphoSitePlus report IMPDH2 expression and candidate phospho-tyrosine residues, functional evidence linking tyrosine phosphorylation to IMPDH2 activity in purine metabolism is currently lacking. To address this, we performed in vitro kinase assays using recombinant human IMPDH2 incubated with catalytically active recombinant ALK or SRC kinases (Fig. 1a). Immunoblotting with pan-phosphotyrosine (pY100) and IMPDH2 antibodies revealed strong tyrosine phosphorylation of IMPDH2 by both kinases (Fig. 1b), which was quantified by densitometric analysis (Fig. 1c). To identify specific phosphorylation sites, IMPDH2 samples from these kinase reactions were analyzed by SDS–PAGE followed by mass spectrometry (Fig. 1d). SRC phosphorylated five residues— Y110, Y233, Y348, Y430, and Y484. At the same time, ALK shared four of these sites and uniquely phosphorylated Y294 (Fig. 1e). Notably, Y110 and Y233 localize to the allosteric regulatory CBS domains of IMPDH2, suggesting a potential mechanism for modulating enzyme activity. Comparative sequence analysis revealed that Y110 in IMPDH2 is substituted for a conserved phenylalanine (F110) in IMPDH1, indicating a functional divergence between isoforms (Supplementary Fig. 3a, b). A schematic overview of the identified phosphorylation sites across IMPDH2 domains is presented in Fig. 1f.

**Fig. 1.**
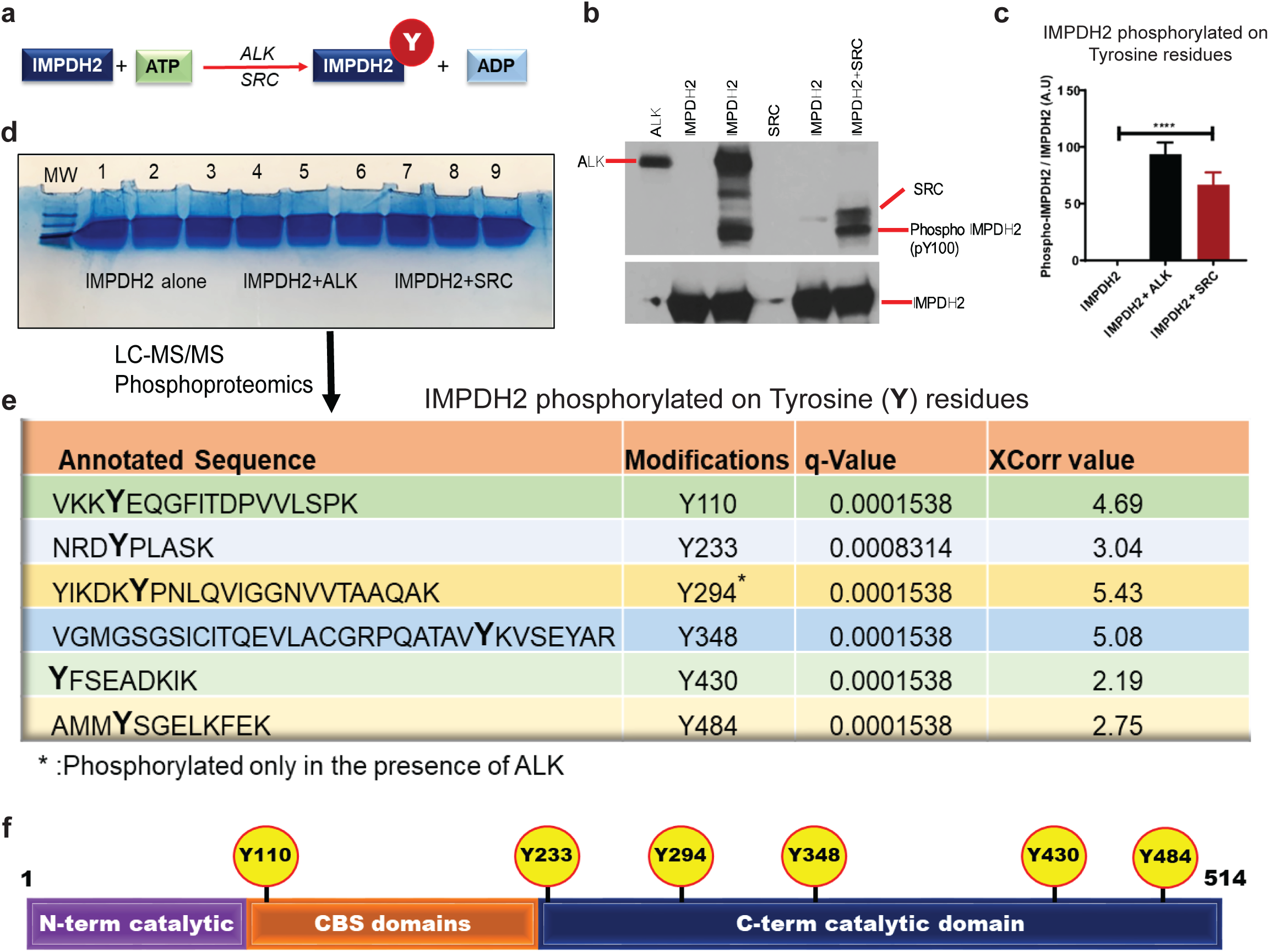
Oncogenic kinases ALK and SRC phosphorylate IMPDH2 at specific tyrosine residues. **a,** Schematic of the in vitro kinase assay used to evaluate tyrosine phosphorylation of IMPDH2 by ALK and SRC kinases. **b,** Western blot analysis of in vitro kinase reactions using ALK and SRC with recombinant IMPDH2, probed with phospho-tyrosine-specific and total IMPDH2 antibodies. **c,** Densitometric quantification of tyrosine-phosphorylated IMPDH2 compared to total IMPDH2 levels, demonstrating enhanced phosphorylation in the presence of ALK and SRC. **d**, Large-scale in vitro kinase assay products were resolved by NuPAGE and analyzed by LC-MS/MS phosphoproteomics to identify phosphorylation sites. **e**, Identification of key tyrosine residues on IMPDH2 phosphorylated by ALK and SRC, as determined by mass spectrometry. **f**, Schematic of IMPDH2 structural domains highlighting novel ALK- and SRC-mediated phosphorylation sites.

### IMPDH2 Y233 phosphorylation regulates PI3P binding

Differences in key lysine (K) and arginine (R) residues between IMPDH1 and IMPDH2 suggest distinct phospholipid binding patterns, warranting further investigation. The most abundant cellular phosphoinositide, PI(4,5)P₂ (PIP2), typically binds to pleckstrin homology (PH) domain proteins, which are critical for BCR signaling and the PI3K/Akt pathways. To examine phospholipid binding, we used PIP strips membranes spotted with various phosphoinositides—to compare recombinant IMPDH1 and IMPDH2 (Fig. 2a). At equal protein concentrations (1 µg/ml), IMPDH1 showed weak binding only to PI3P, while IMPDH2 strongly bound PI3P, PI(4)P, and phosphatidic acid (PA) (Fig. 2b,c). Dilution of IMPDH2 to 100 ng/ml confirmed strong and specific binding to PI3P (Fig. 2d), with densitometric analysis quantifying this interaction (Fig. 2e). To obtain a more quantitative measure, we performed a PIP array assay featuring a gradient of phosphoinositide concentrations (Fig. 2f). This revealed a clear, concentration-dependent binding affinity of IMPDH2 for PI3P (Fig. 2g). Based on our kinase and phosphoproteomic data, we hypothesized that tyrosine phosphorylation at Y233 modulates IMPDH2’s interaction with PI3P. To test this, 293T cells were transiently transfected with IMPDH2-HA alone or with active SRC kinase. Lysates subjected to a PI3P binding assay showed that IMPDH2’s PI3P binding was significantly reduced in the presence of SRC kinase (Fig. 2h). Finally, co-transfection of IMPDH2-mCherry with the PI3P marker EGFP-FYVE in 293T cells revealed strong co-localization by microscopy, confirming IMPDH2’s cellular association with PI3P-enriched membranes (Fig. 2i).

**Fig. 2.**
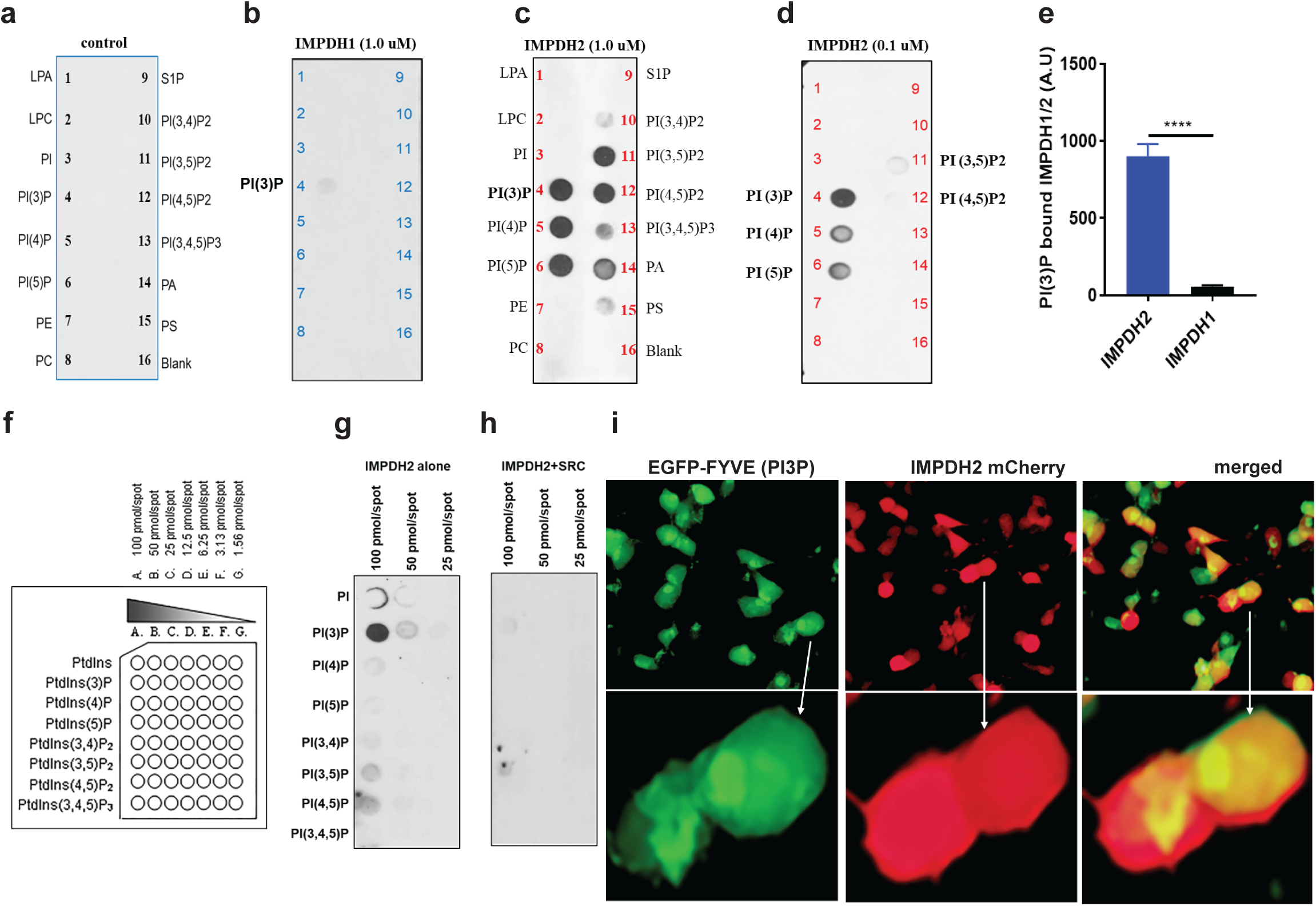
IMPDH2 Y233 phosphorylation regulates PI3P binding. **a,** Schematic of PIP-lipid strip assay showing membranes coated with 15 distinct phospholipid species and incubated with HA peptide as a negative control. **b**, Recombinant human IMPDH1 (1.0 µg) incubated on PIP-lipid strips in 3% BSA/PBS overnight at 4°C; binding detected by Western blot using IMPDH1 antibody. **c**, Recombinant human IMPDH2 (1.0 µg) incubated under identical conditions and probed with IMPDH2 antibody. **d**, Control binding experiment using a reduced amount (0.1 µg) of IMPDH1 and probing with IMPDH2 antibody to assess specificity. **e**, Densitometric quantification of PI3P binding to IMPDH2 and IMPDH1, demonstrating preferential binding of PI3P to IMPDH2. **f**, Schematic of membrane-lipid arrays containing eight phosphoinositide species used to assess phosphoinositide-specific interactions. **g, h**, HA-tagged human IMPDH2 constructs expressed in HEK293T cells, with or without co-expression of SRC kinase; PI3P-bound IMPDH2-HA was detected by Western blot using anti-HA antibody. **i,** Live-cell imaging of HEK293T cells co-transfected with IMPDH2-mCherry and EGFP-FYVE (PI3P biosensor), revealing co-localization of IMPDH2 with PI3P in the presence of Y233 phosphorylation. All graphs represent mean ± SD from three biological replicates (n = 3). Statistical significance was assessed using unpaired t-tests: *p < 0.05, **p < 0.01, ***p < 0.001, ****p < 0.0001.

### PI3P Binding Inhibits IMPDH2 Activity

Protein–phospholipid interactions often regulate protein function. To assess the effect of PI3P on IMPDH2 enzymatic activity, we performed activity assays using purified recombinant IMPDH2 in the presence of synthetic PI3P and PIP2 (Fig. 3a, b). PI3P caused a dose-dependent inhibition of IMPDH2 activity, with 200 µM reducing enzyme activity by over 50% (p < 0.001) (Fig. 3c). In contrast, IMPDH1 activity was unaffected by PI3P (Fig. 3d). Using mycophenolic acid (MPA), a known inhibitor of both IMPDH1 and IMPDH2, as a positive control, we confirmed that MPA inhibited both isoforms, whereas PI3P selectively inhibited IMPDH2 (Fig. 3e). PIP2 treatment did not alter the activity of either enzyme (Fig. 3f, g). These results demonstrate that PI3P specifically and dose-dependently inhibits IMPDH2 activity without affecting IMPDH1, while MPA inhibits both enzymes, and PIP2 has no effect.

**Fig. 3.**
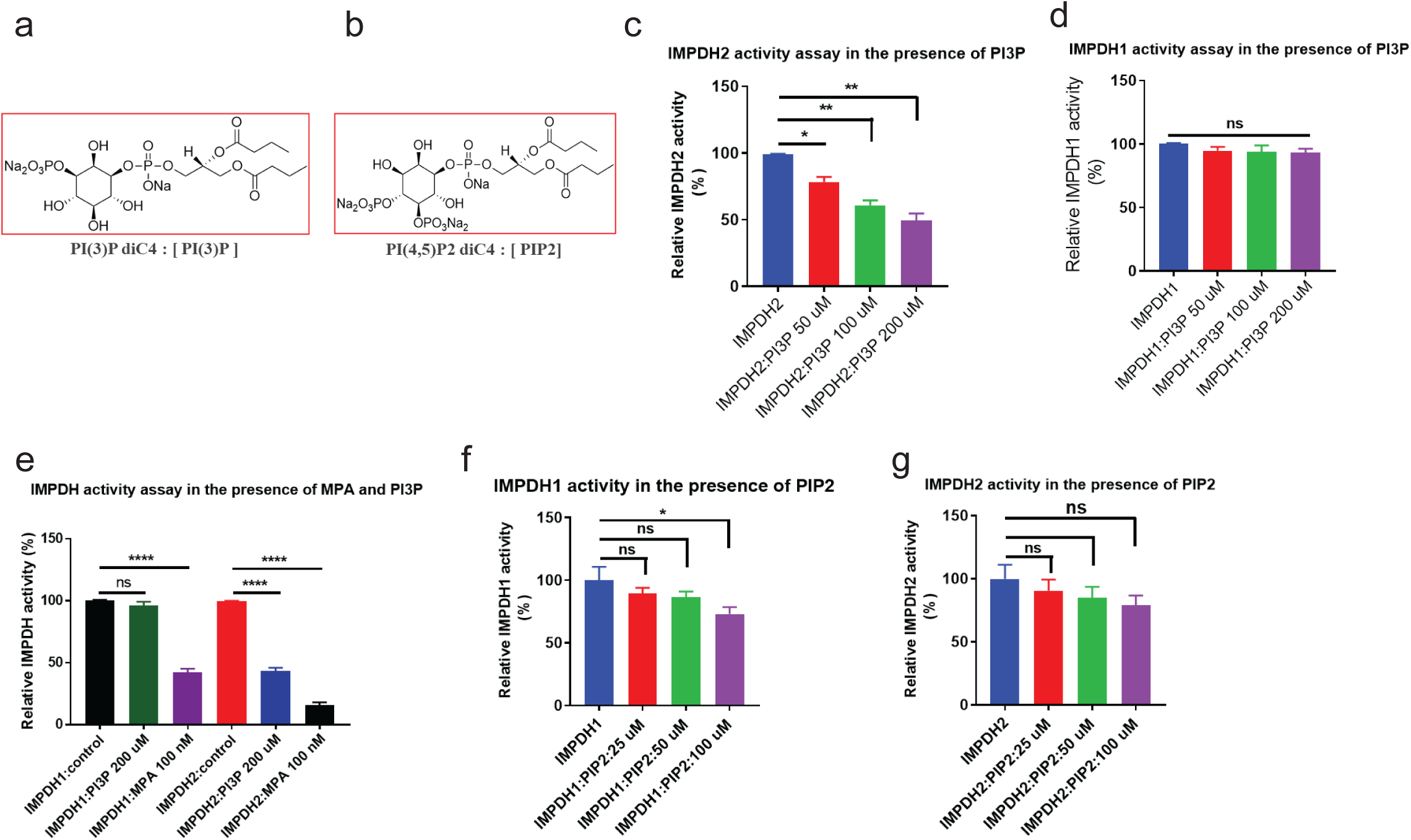
PI3P binding inhibits IMPDH2 enzymatic activity. **a, b**, Chemical structures of phospholipids PI3P and PIP2 used in this study. **c**, Recombinant IMPDH2 was pre-incubated with increasing concentrations of synthetic PI3P; enzymatic activity was measured and showed dose-dependent inhibition. **d**, Recombinant IMPDH1 was similarly pre-incubated with PI3P to assess its sensitivity to PI3P-mediated inhibition. **e**, Comparative enzymatic activity of recombinant IMPDH1 and IMPDH2 pre-incubated with 200 µg synthetic PI3P or MPA (mycophenolic acid, a known IMPDH inhibitor) as a positive control. **f**, Recombinant IMPDH1 was pre-incubated with increasing concentrations of synthetic PIP2; no significant inhibition of enzymatic activity was observed. **g**, Recombinant IMPDH2 pre-incubated with synthetic PI3P confirms reproducible, dose-dependent inhibition of enzymatic activity. All graphs display mean ± SD from three biological replicates (n = 3). Statistical significance was calculated using unpaired t-tests: *p < 0.05, **p < 0.01, ***p < 0.001, ****p < 0.0001.

### IMPDH1-derived synthetic peptide acts as an inhibitor of IMPDH2

Previous studies have emphasized the critical role of peptide–protein interactions (PPIs) in cellular signaling and regulatory networks(29, 30). IMPDH enzymes are catalytically active in solution and typically crystallize as tetramers or octamers(31, 32). The CBS domain is key to IMPDH allosteric regulation. We hypothesized that self-inhibitory peptides from CBS domains could interfere with enzyme activity. Given 85% sequence similarity between IMPDH1 and IMPDH2, we synthesized two peptides targeting critical CBS regions (Fig. 4a). The Y233 residue, highly conserved across species (Fig. 4b, c), was central to our focus since ALK kinase phosphorylation of IMPDH2 may alter its allosteric regulation by reducing PI3P binding. We designed peptides from the N-terminal CBS domain (aa 122–136), rich in lysine and arginine in IMPDH2, and from the C-terminal CBS domain (aa 228–242), which includes Y233 and a Y233F mutant (Fig. 4d, e). Enzymatic assays showed no significant change in IMPDH1 activity with CBS-1 peptides (Fig. 4f), but IMPDH2 activity significantly decreased over >43% with the IMPDH2 CBS-1 peptide (Fig. 4g). Using CBS-2 peptides, IMPDH1 activity dropped >80% with its own Y233 peptide but not with Y233F (Fig. 4h). IMPDH2 CBS-2 peptides had no effect on IMPDH1 unless Y233 was mutated (Fig. 4i). IMPDH2 activity reduced >91% with IMPDH1 CBS-2 Y233 peptide versus ∼10% with Y233F (Fig. 4j). In contrast, self-derived CBS-2 peptides inhibited IMPDH2 significantly (Fig. 4k). Since CBS-2 mediates nucleotide binding, we tested AMP, ATP, GMP, GDP, and GTP. AMP and ATP moderately inhibited IMPDH2, while GTP caused potent inhibition (Fig. 4l, m). We obtained the 3D structure of the 15-mer peptide (aa 228–242) from AlphaFold and performed energy minimization with AMBER-EHT. Using hydrophobic patch analysis, we docked the peptide onto IMPDH2, identifying 100 binding sites and selecting the top five based on energy scores (Fig. 4n). The peptide binds between the catalytic and GTP-binding domains, thereby restricting the function of IMPDH2. Binding specificity involves hydrogen bonds between the peptide’s tyrosine and IMPDH2 residues Y233 and H454 (highlighted in yellow). In IMPDH1, Q454 replaces H454 (data not shown). Substituting Y with F disrupts these interactions, preventing peptide binding and preserving enzyme activity. Thus, molecular docking reveals that CBS domain peptides inhibit IMPDH2 by binding near the GTP site, with Y233 critical for binding and regulation.

**Fig. 4.**
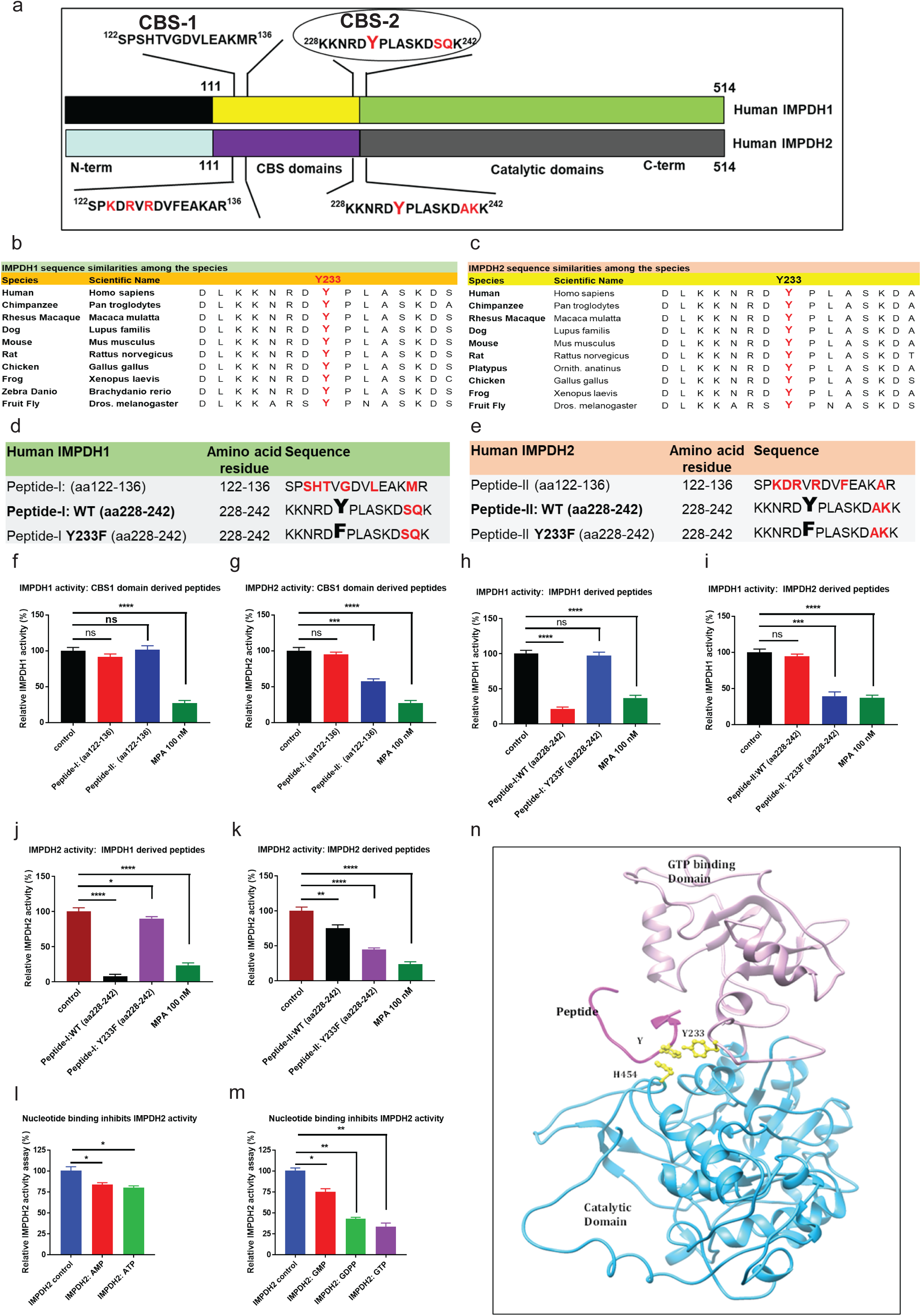
IMPDH1-derived synthetic peptide inhibits IMPDH2 enzymatic activity. **a**, Schematic representation of human IMPDH1 and IMPDH2 structural domains highlighting CBS-1 and CBS-2 regions, which contribute to allosteric regulation. The Y233-containing peptide—central to this study—is located near the CBS-2 domain and is phosphorylated by ALK and SRC kinases. **b, c**, Sequence alignment of human IMPDH1 and IMPDH2 showing conservation of the Y233 region across species, underscoring its potential functional importance. **d, e**, Sequences of synthetic peptides derived from CBS-1 and CBS-2 domains of IMPDH1 and IMPDH2 used in enzymatic activity assays. **f**, IMPDH1 activity assay in the presence of CBS-1-derived peptides from IMPDH1 and IMPDH2, with MPA included as a positive control for inhibition. **g,** IMPDH2 activity assay under the same conditions as in (f), assessing peptide-mediated inhibition. **h**, IMPDH1 activity assay with CBS-2 domain peptides from IMPDH1, comparing wild-type (Y233) versus Y233F mutant (tyrosine to phenylalanine substitution). **i**, IMPDH1 activity in the presence of CBS-2 peptides from IMPDH2, comparing WT vs. Y233F variant. **j**, IMPDH2 activity assay in the presence of CBS-2 peptides from IMPDH1 (WT and Y233F). **k**, IMPDH2 activity in the presence of CBS-2 peptides from IMPDH2 (WT and Y233F), showing that phosphorylation state of Y233 affects inhibition. **l**, IMPDH2 enzymatic activity following pre-incubation with AMP or ATP, testing nucleotide-based allosteric regulation. **m**, IMPDH2 enzymatic activity in response to GMP, GDP, and GTP, to assess guanine nucleotide-mediated feedback. **n**, Ribbon model of IMPDH2 showing bound inhibitory peptide. The catalytic domain is shown in cyan, GTP-binding domain in light pink, and the peptide (15-mer) in magenta. Three peptide-interacting residues are highlighted in yellow, indicating key contact points. All graphs represent mean ± SD from three biological replicates (n = 3). Statistical analyses were performed using unpaired t-tests: *p < 0.05, **p < 0.01, ***p < 0.001, ****p < 0.0001.

### *Structure-based in silico* screening and discovery of a novel allosteric inhibitor targeting IMPDH2

Building on our findings that IMPDH2 Y233 phosphorylation, PI3P binding, peptide-based inhibition, and GTP all regulate IMPDH2 activity via the allosteric domain, we identified lead compound-10 as a novel and selective IMPDH2 inhibitor. For in silico screening, we used the published PDB structure of the IMPDH2 allosteric GTP-binding domain(33) as shown (Fig. 5a, b). We performed in silico screening using the National Cancer Institute (NCI) Mechanistic Set VI, comprising 811 compounds selected from 37,836 based on diverse growth inhibition profiles in the NCI-60 cell line screen. Compounds were clustered using the FASTCLUS algorithm in SAS, yielding 1,272 clusters, from which representative compounds were selected. After applying filters (MW 100–500, single molecule, 2–6 rotatable bonds, Lipinski drug-like = 1, no chiral centers), 283 compounds remained. Approximately 15,200 energetically favorable conformations were generated and docked into the GTP-binding site of IMPDH2. The top 100 compound-IMPDH2 complexes were ranked, and 38 unique compounds were identified based on NSC identity (Fig. 5c). We obtained 15 compounds from the NCI, and their 2D structures are shown (Fig. 5d). From these, 25 structural analogs were also identified (Supplementary Fig. 4). Initial enzyme activity screening at 10 µM identified three active compounds: comp-5 (>32% inhibition), comp-10 (>99%), and comp-12 (>52%). At 1 µM, comp-10 showed the most potent inhibition (>93%) of recombinant human IMPDH2 activity. Docking analysis of comp-10 with IMPDH2 is shown (Fig. 5e). Dose-response assays comparing comp-10 and mycophenolic acid (MPA) yielded IC₅₀ values of ∼260 nM and ∼106 nM, respectively (Fig. 5f, g).

**Fig. 5.**
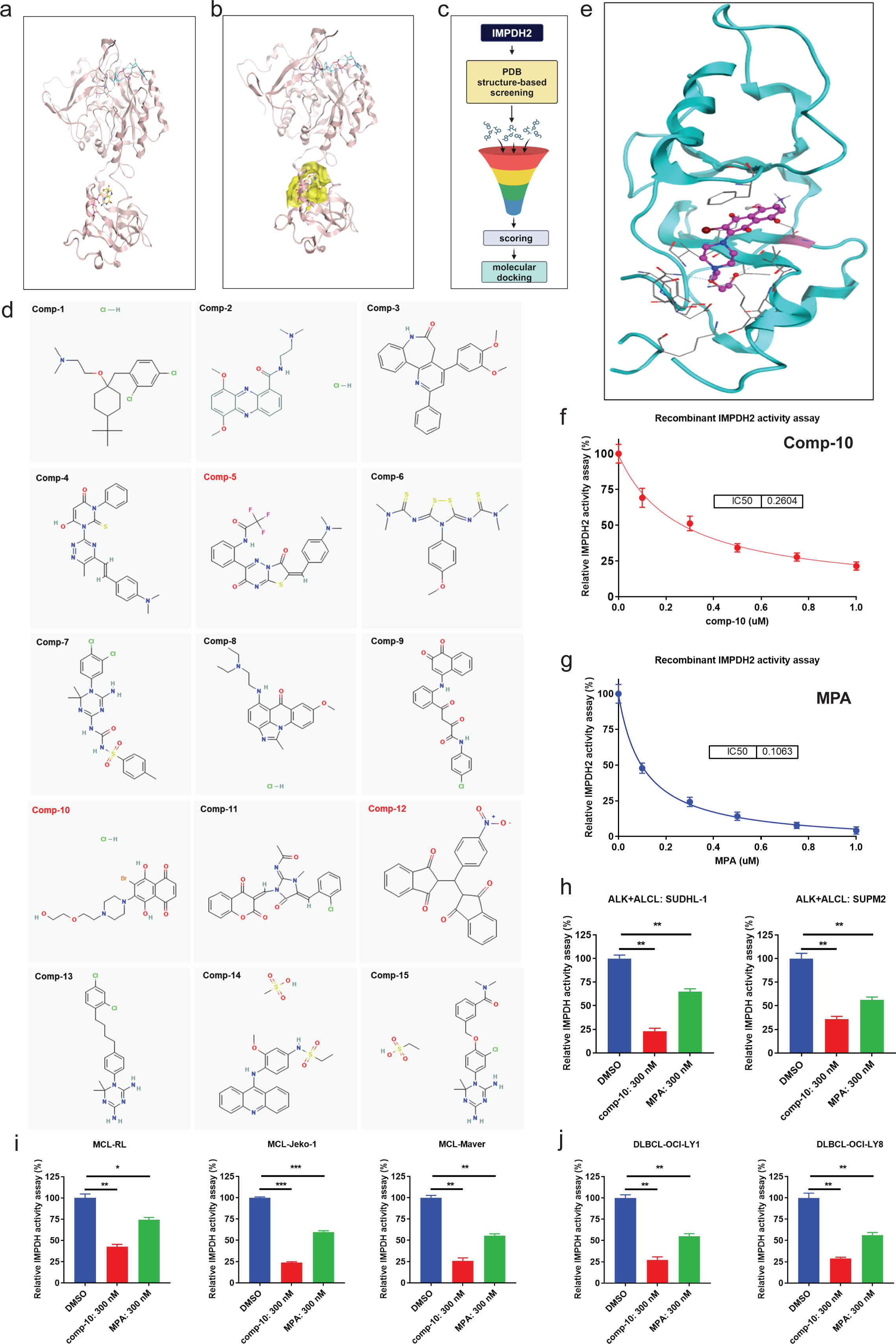
Structure-based virtual screening identifies a novel small-molecule inhibitor of human IMPDH2. **a**, Ribbon diagram of the CBS (cystathionine β-synthase) domain of human inosine monophosphate dehydrogenase 2 (IMPDH2), illustrating the secondary structural elements. **b**, Surface representation of the GTP-binding pocket (highlighted in yellow) within the IMPDH2 structure, showing the targeted allosteric site for compound docking. **c**, Schematic overview of the in silico screening pipeline, including structure-based virtual screening, docking, and filtering steps leading to hit identification. **d**, Chemical structures of 15 top-ranked compounds identified from virtual screening; compound 10 (comp-10) was selected for further validation based on docking scores and predicted binding interactions. **e**, Molecular docking pose of comp-10 within the GTP-binding pocket of human IMPDH2 (based on the PDB reference structure), revealing key interactions with the binding site residues. **f**, In vitro enzymatic activity assay of recombinant IMPDH2 in the presence of comp-10, showing concentration-dependent inhibition and determination of the half-maximal inhibitory concentration (IC₅₀). **g**, Enzymatic assay using the clinically established IMPDH inhibitor mycophenolic acid (MPA) as a positive control for benchmarking IC₅₀ and assay reproducibility. **h–j**, Inhibition of endogenous IMPDH activity by comp-10 in human lymphoma cell lines: **h**, anaplastic large cell lymphoma (ALCL); **i,** mantle cell lymphoma (MCL); and **j,** diffuse large B-cell lymphoma (DLBCL). Comp-10 treatment significantly reduced IMPDH enzymatic activity across all tested cell lines. All the graphs show mean ± SD (n = 3 biological replicates), and all statistical analyses were conducted with unpaired t-tests: ^∗^p < 0.05, ^∗∗^p < 0.01, ^∗∗∗^p < 0.001, ^∗∗∗∗^p < 0.0001.

Additionally, in silico ADME profiling of Comp-10 was performed using the SwissADME web tool to assess its pharmacokinetics, drug-likeness, and medicinal chemistry friendliness, as described previously(34). To compare ADME (Absorption, Distribution, Metabolism, and Excretion) properties, we included mycophenolic acid (MPA), a known inhibitor of IMPDH, as a positive control. SwissADME analysis showed that Comp-10 exhibited similar ADME properties to MPA across all pharmacokinetic and drug-likeness parameters (Supplementary Fig. 5a, b) and fully adhered to Lipinski’s Rule of Five. Based on the Comp-10 structure, we have also identified several analogs that are currently being tested for IMPDH inhibition and cell growth suppression. To assess cellular IMPDH1/2 inhibition, we treated ALCL (SUDH-L1, SUPM2) and MCL (RL, Jeko-1, Maver) cell lines with 300 nM Comp-10 for 24 hours. IMPDH activity assays using equal amounts of DMSO- and Comp-10-treated lysates (5–10 µg per well, N = 3) revealed strong inhibition in SUDH-L1 (>77%; p < 0.001) and SUPM2 (>64%; p < 0.001), compared to modest inhibition by MPA (36% and 44%, respectively; Fig. 5h). Similarly, Comp-10 significantly reduced IMPDH activity in MCL lines: RL (>58%; p < 0.001), Jeko-1 (>74%; p < 0.0001), and Maver (>75%; p < 0.0001) (Fig. 5i). In DLBCL cell lines OCI-LY1 and OCI-LY8, Comp-10 led to robust inhibition exceeding 73% and 71%, respectively (p < 0.001) (Fig. 5j).

### Comp-10 inhibits the expression of IMPDH1 and IMPDH2 and does not induce the formation of IMPDH filaments

We compared the effects of Comp-10 and MPA on IMPDH1 and IMPDH2 expression in ALCL cell lines, both sensitive and resistant to ALK inhibitors (ALKi) crizotinib and lorlatinib. Treatment of ALKi-sensitive SUPM2 and L82 cells with 300 nM Comp-10 for 48 hours led to a marked reduction in both IMPDH1 and IMPDH2 protein levels, whereas MPA modestly increased their expression (Fig. 6a). Densitometric analysis of IMPDH2 and IMPDH1 shows a significant reduction in Comp-10 treated samples (Fig.6b, c). Similar results were observed in crizotinib- and lorlatinib-resistant ALCL cell lines (Karpas299 and SUPM2-CR), where Comp-10 reduced IMPDH1/2 levels, while MPA increased them (Fig. 6d–f). Next, we tested comp-10, MPA, and MF on the MCL cell line Maver. Quantitative RT-PCR was performed to determine the expression levels of IMPDH2. The results showed that Comp-10 did not induce an increase in IMPDH2 expression, whereas both MPA and MF significantly upregulated IMPDH2 expression (Fig.6g). Furthermore, Comp-10 and MPA differentially regulate IMPDH1 and IMPDH2 expression in BTKi-sensitive and -resistant MCL cell lines. Western blot analysis revealed that treatment with Comp-10 (300 nM) led to a marked reduction in IMPDH1 and IMPDH2 protein levels across all tested MCL cell lines, while MPA treatment resulted in a significant upregulation of both proteins. In Maver cells, qRT-PCR showed that Comp-10 did not alter IMPDH2 mRNA levels, whereas MPA and MF significantly increased IMPDH2 transcription. Expression of MYC, a known transcriptional regulator of IMPDH1/2, remained unchanged, suggesting that Comp-10 regulates IMPDH1/2 post-transcriptionally (Fig. 6h). Densitometric quantification confirmed a significant decrease in IMPDH1 and IMPDH2 protein levels following Comp-10 treatment (Fig. 6i). We also evaluated IMPDH filament formation (rods and rings) in SUDHL-1 cells expressing HA-GFP-tagged IMPDH2. DMSO-treated cells showed no filaments (Fig. 6j), MPA induced prominent rod/ring formation (Fig. 6k), while Comp-10-treated cells did not exhibit filament formation (Fig. 6l). In summary, IMPDH2 is overexpressed relative to IMPDH1 in ALCL and MCL cell lines. The catalytic-site inhibitor MPA upregulated IMPDH1/2 and promoted rod/ring formation— potentially linked to toxicity. In contrast, Comp-10, which targets the regulatory domain, downregulated IMPDH1/2, blocked filament formation, and was effective in both ALKi-sensitive and -resistant models. Importantly, Comp-10 acted post-transcriptionally, offering a mechanistically distinct and potentially superior alternative to existing IMPDH inhibitors.

**Fig. 6.**
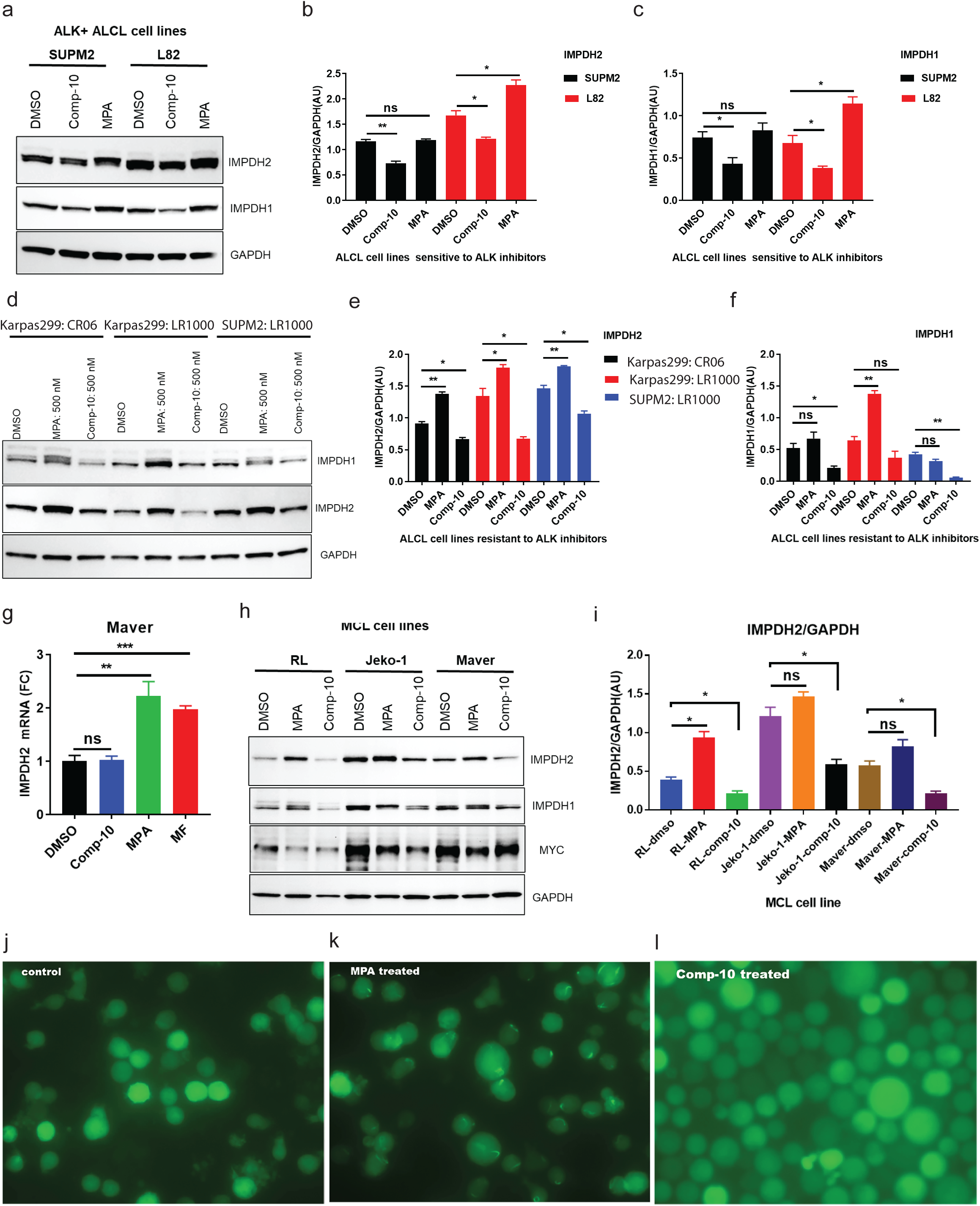
Comp-10 downregulates IMPDH1/2 expression and prevents filament (rod/ring) formation in ALCL and MCL models. **a**, Western blot analysis of IMPDH1 and IMPDH2 expression in ALK inhibitor (ALKi)-sensitive ALCL cell lines (SUPM2 and L82) following 48-hour treatment with Comp-10 (300 nM) or mycophenolic acid (MPA). **b, c**, Densitometric quantification of IMPDH1 and IMPDH2 protein bands from panel **a**, confirming significant downregulation following Comp-10 treatment. **d**, Western blot of ALKi-resistant ALCL cell lines (Karpas299 and SUPM2-CR) treated with Comp-10 and MPA, showing consistent downregulation of IMPDH1/2 by Comp-10 and upregulation by MPA. **e, f**, Densitometric analysis of protein expression in resistant ALCL lines (panel **d**) supports differential regulation by Comp-10 and MPA. **g**, Quantitative RT–PCR analysis of IMPDH2 mRNA expression in MCL cell line Maver treated with Comp-10, MPA and MF. **h**, Western blot analysis of BTK inhibitor (BTKi) sensitive and resistant MCL cell lines treated with Comp-10 and MPA. **i**, Densitometric quantification of Western blots shown in panel **h**, confirming significant downregulation of IMPDH1/2 protein levels by Comp-10 across multiple MCL models. **j–l**, Light microscopy of SUDHL-1 cells expressing HA–GFP–IMPDH2 shows distinct patterns of filament formation: DMSO-treated control cells (**j**) lack filaments; MPA-treated cells (**k**) display prominent rod/ring structures; and Comp-10-treated cells (**l**) show no rod/ring or filament formation. Densitometric values of the Western blots (N=2). All the graphs show mean ± SD (n = 3 biological replicates), and all statistical analyses were conducted with unpaired t-tests: ^∗^p < 0.05, ^∗∗^p < 0.01, ^∗∗∗^p < 0.001, ^∗∗∗∗^p < 0.0001.

### Comp-10 inhibits the growth and colony formation of malignant ALK-positive cells

To evaluate the effect of comp-10 on the growth of malignant cells, we focused on ALCL cell lines as a cancer model, using MPA as a positive control. While comp-10 and MPA displayed a similar inhibitory effect on one of the lines (Fig. 7a), comp-10 showed a more substantial inhibitory effect than MPA in two other lines (Fig. 7b,c). Notably, it potently inhibited the growth of not only the parental Karpas 299 cell line, which is sensitive to ALK inhibitors (Fig. 7d), but also its derivative cell lines resistant to ALK inhibitors Crizotinib (Fig. 7e) and Lorlatinib (Fig. 7f). Similar results were obtained with the sensitive parental SUP-M2 (Fig. 7g) as well as its Crizotinib- and Lorlatinib-resistant derivatives (Fig. 7h, i). Next, we tested the effect of Comp-10 on colony formation. Here, we used a stable SUDHL-1 cell line tagged with IMPDH2-HA-GFP. Briefly, 200 cells per well in triplicate (in 6-well plates) were plated in Human Methylcellulose media, following the manufacturer’s protocol, with a DMSO control, comp-10 (100 nM), and MPA (100 nM). After plating, the plates were incubated in a standard cell culture chamber for four weeks. Colonies were then photographed using the iBright imaging system in GFP fluorescence mode (Fig. 7j). Cell counts were performed, and the results showed that cells treated with comp-10 formed significantly fewer colonies (>12; p < 0.0001) compared to the DMSO control. Additionally, MPA treatment resulted in more than 37 colonies (p < 0.002) relative to DMSO (Fig. 7k). Colony numbers were expressed as a percentage of the DMSO control, indicating that comp-10–treated cells formed significantly fewer colonies (>12%), while MPA-treated cells formed over 72% compared to DMSO (Fig. 7l). Next, we tested comp-10 on two index cell lines from MCL and found that comp-10 suppresses cell growth more effectively than MPA. (Fig. 7m, n). Notably, the growth of the OVCAR3 ovarian carcinoma cell line was unaffected by either comp-10 or MPA (Fig. 7o), indicating a lack of nonspecific toxicity from both MPA and comp-10.

**Fig. 7.**
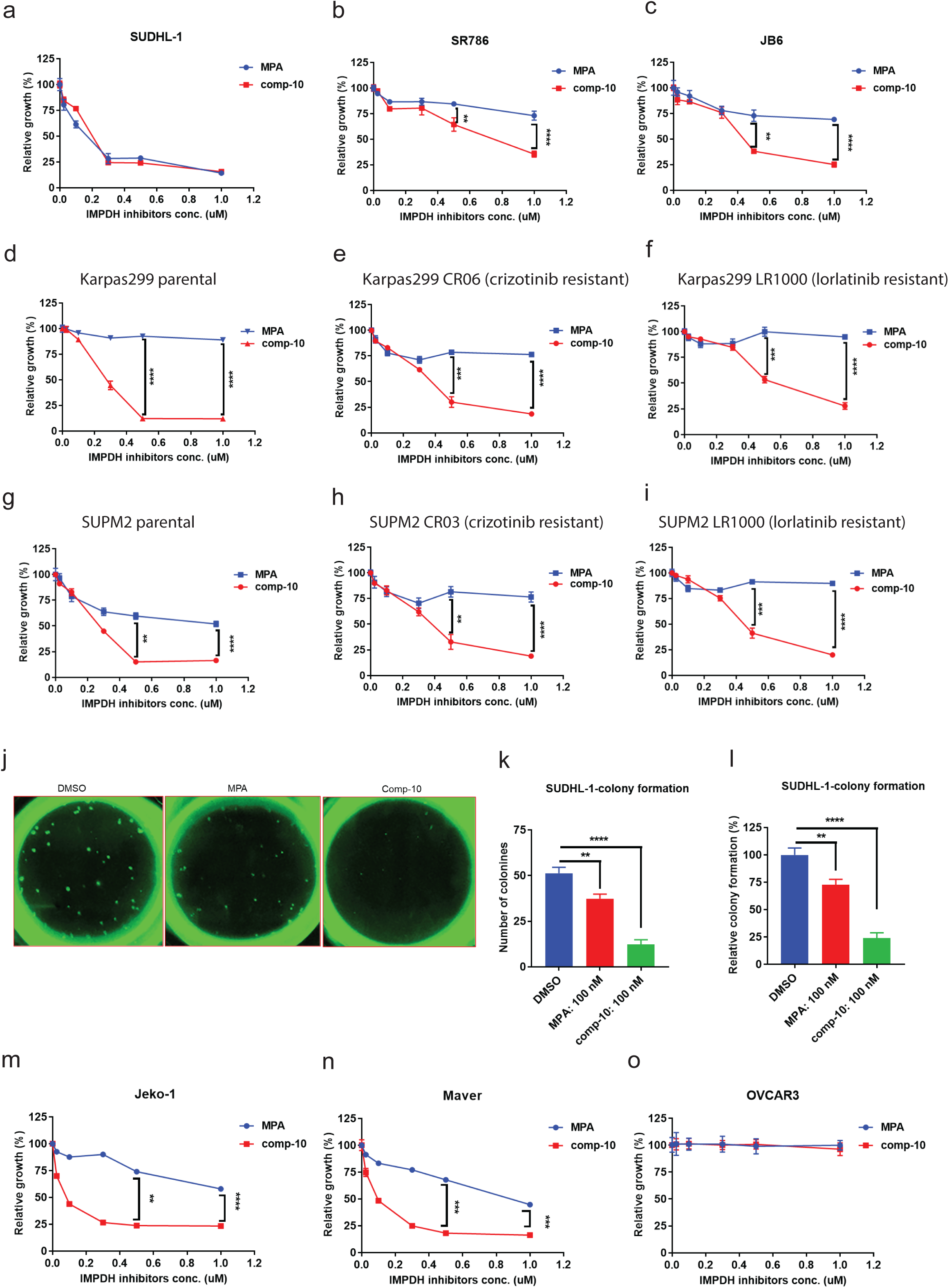
Comp-10 inhibits proliferation and colony formation in ALK-positive lymphoma cells. **a–c**, Cell viability assays of ALCL cell lines treated with comp-10 or mycophenolic acid (MPA). While both compounds similarly inhibited growth in one line (**a**), comp-10 showed significantly stronger growth inhibition in two additional lines (**b**, **c**). **d–f**, Comp-10 potently suppressed the growth of ALK inhibitor (ALKi)-sensitive Karpas 299 cells (**d**) and their resistant derivatives: Crizotinib-resistant (**e**) and Lorlatinib-resistant (**f**) lines. **g–i**, Similar results were observed in parental SUP-M2 cells (**g**) and their ALKi-resistant derivatives (**h**, **i**), with comp-10 maintaining robust antiproliferative activity. **j**, Representative GFP fluorescence images of colonies formed by SUDHL-1 cells stably expressing IMPDH2–HA–GFP in methylcellulose after 4 weeks of treatment with DMSO, comp-10 (100 nM), or MPA (100 nM). **k**, Quantification of colony numbers per well. Comp-10 significantly reduced colony formation compared to both DMSO (p < 0.0001) and MPA (p < 0.01). **l**, Colony numbers expressed as percentage of DMSO control. Comp-10 reduced colony formation to ∼12%, while MPA-treated cells retained >72% colony formation relative to control. **m, n**, Cell viability assays in two mantle cell lymphoma (MCL) index lines demonstrate that comp-10 effectively inhibits cell growth than MPA. **o**, Growth of the ovarian carcinoma cell line OVCAR3 remained unaffected by comp-10 or MPA, indicating minimal off-target toxicity in non-lymphoid malignancies. All the graphs show mean ± SD (n = 3 biological replicates), and all statistical analyses were conducted with unpaired t-tests: ^∗^p < 0.05, ^∗∗^p < 0.01, ^∗∗∗^p < 0.001, ^∗∗∗∗^p < 0.0001.

**Fig. 8.**
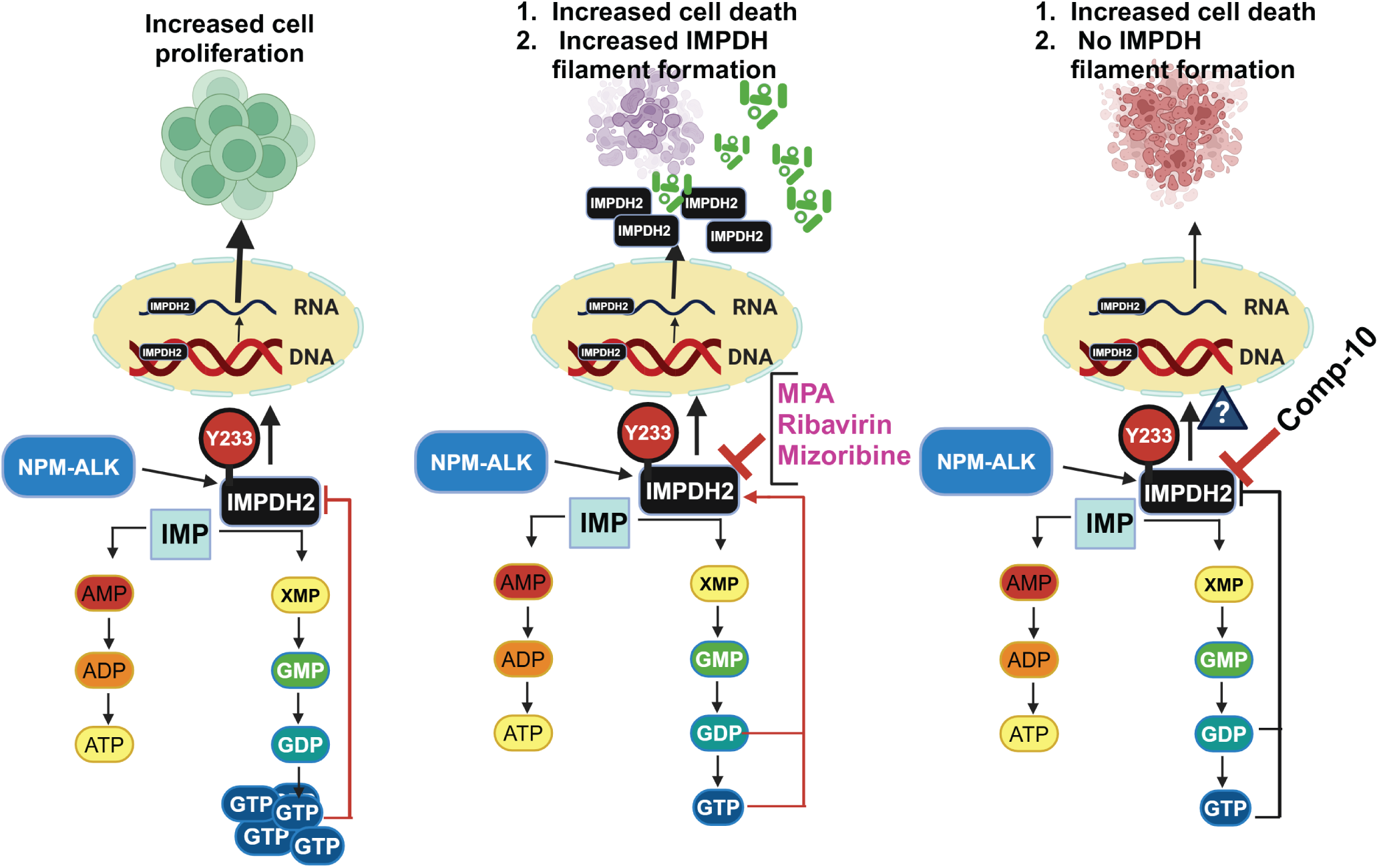
Graphical summary of the discovery. Tyrosine phosphorylation of IMPDH2 enhances its activity, contributing to increased cell proliferation. Traditional IMPDH inhibitors (e.g., MPA) block enzymatic function but disrupt GTP-mediated feedback inhibition, often triggering the formation of IMPDH filaments (rods and rings), which is associated with cellular stress and toxicity. In contrast, Comp-10 inhibits IMPDH activity while preserving regulatory control and preventing filament formation, offering a distinct and potentially less toxic mechanism of action.

## Discussion

Our study provides the first direct evidence that oncogenic tyrosine kinases (TKs), specifically ALK and SRC, phosphorylate IMPDH2, a critical metabolic enzyme. TKs are well-established drivers of cancer progression and patient outcomes, largely through modulating cellular signaling networks and metabolic pathways via post-translational modifications. While tyrosine phosphorylation comprises a small fraction of total protein phosphorylation, its influence on cell physiology, including metabolism, survival, and proliferation, is profound. There are five FDA-approved ALK inhibitors for EML4-ALK-positive non-small cell lung cancer (NSCLC): crizotinib, ceritinib, alectinib, brigatinib, and lorlatinib. Notably, crizotinib is also approved for treating NPM-ALK-positive anaplastic large cell lymphoma (ALCL) and neuroblastoma. Despite these advancements, many patients still experience resistance to all these ALK inhibitors, emphasizing the need for continued research and the development of new therapeutic approaches. Although IMPDH2 overexpression has been widely observed in cancers, the functional role of its post-translational modifications, especially tyrosine phosphorylation, remained largely unexplored. Our findings reveal that phosphorylation at the conserved Y233 residue within the IMPDH2 allosteric domain enhances its enzymatic activity, linking TK signaling directly to metabolic reprogramming in cancer cells. This mechanistic insight has significant therapeutic implications. Targeting key downstream substrates of oncogenic TKs, such as IMPDH2 may circumvent resistance mechanisms that limit the efficacy of direct TK inhibitors. Our demonstration that IMPDH inhibition suppresses the growth of both ALK inhibitor-sensitive and -resistant ALCL cells positions IMPDH2 as a compelling therapeutic target. Furthermore, since other TKs such as SRC are involved in various cancers, this approach could be applicable more broadly.

Building on our identification of PI3P as a natural inhibitor that binds IMPDH2 and modulates its function, we designed a peptide centered on the Y233 site. PI3P’s role in membrane trafficking and autophagy, combined with its specific binding to basic residues within the CBS domains of IMPDH2, appears to disrupt tetramer formation and enzyme activity. PI3P plays a crucial role in membrane dynamics and trafficking because it is linked to endosomal membranes and autophagy function(35). The active site of IMPDH2, Cys-331, is located in the catalytic domain, which differs from the site of IMPDH, Y233. The active site Cys-331 forms a covalent thio-adduct with the typical substrate IMP in the IMPDH—MPA–IMP complex(31). IMPDH2 Y233 is located within the allosteric and GTP-binding domains, highlighting its crucial role in allosteric regulation that warrants further investigation. These findings highlight the potential for allosteric modulation of IMPDH2 as a novel therapeutic avenue distinct from classical active-site inhibition.

The drug discovery process remains challenging and costly despite significant advances in basic life sciences and biotechnology. It takes about 10 to 15 years and roughly $2 billion for a small-molecule inhibitor to move from the laboratory to patient care. Meanwhile, computational approaches are streamlining drug discovery through structure-based virtual screening of large chemical spaces, significantly improving the ability to find hit targets while reducing costs and time(36). In this context, our computational drug discovery approach led to the identification of Comp-10, a first-in-class allosteric inhibitor of IMPDH. Unlike traditional inhibitors such as mycophenolic acid (MPA), which induce polymerization of IMPDH filaments (rods and rings) and paradoxically increase IMPDH expression, Comp-10 uniquely decreases IMPDH protein levels and does not promote filament formation. This novel mechanism may improve therapeutic efficacy and reduce resistance. In summary, our work elucidates a direct regulatory axis between oncogenic TKs and IMPDH2, mediated by phosphorylation at Y233, which sustains the metabolic demands of cancer cells. Targeting this axis with allosteric inhibitors, such as Comp-10, offers a promising strategy to overcome resistance and improve outcomes in ALK-driven and potentially other TK-driven malignancies. Future studies will explore the translational potential of these findings and assess the broader applicability of allosteric IMPDH inhibitors in cancer and other diseases characterized by dysregulated metabolism. These results validate comp-10 as a promising lead compound for targeting IMPDH2. The findings suggest that inhibiting IMPDH with the novel compound comp-10 provides clear therapeutic benefits over MPA.

In summary, we found that TK, represented by ALK and SRC kinase, phosphorylates IMPDH2 at a highly conserved Y233 site within the allosteric domain, fostering the enzymatic activity of this key metabolic enzyme. The interaction between IMPDH2 and PI3P was identified as an inhibitor of IMPDH2, and synthetic peptides derived from IMPDH1 were found to suppress the enzymatic activity of IMPDH2. Comp-10, the first-in-class allosteric IMPDH inhibitor, inhibited the growth of malignant ALCL and MCL cells. These findings suggest that allosteric IMPDH inhibitors could offer a novel therapeutic approach to cancer and possibly other diseases.

## Acknowledgments

The Mass Spectrometry and Proteomics Core at the University of Pennsylvania’s School of Medicine, Philadelphia, PA, performed the phosphoproteomics analysis. We have created the figures in this manuscript using BioRender program.

## Author Contributions

J.B. conceived the project, designed and performed the research, analyzed the data, supervised the project, and wrote the original draft and edited the final draft; C.L. performed the experiments and reviewed the manuscript; D.R. performed experiments; L.M. provided the ALK inhibitor-resistant cell lines and reviewed the manuscript; N.P. performed in-silico screening and molecular modeling, acquired the compounds from the NCI repository, wrote the in-silico drug discovery aspects, and reviewed the manuscript; M.A.W. provided resources, reviewed and edited the manuscript, and obtained the funding for the project.

## Figure legends for the supplementary Figures

**Supplementary Fig. 1.**
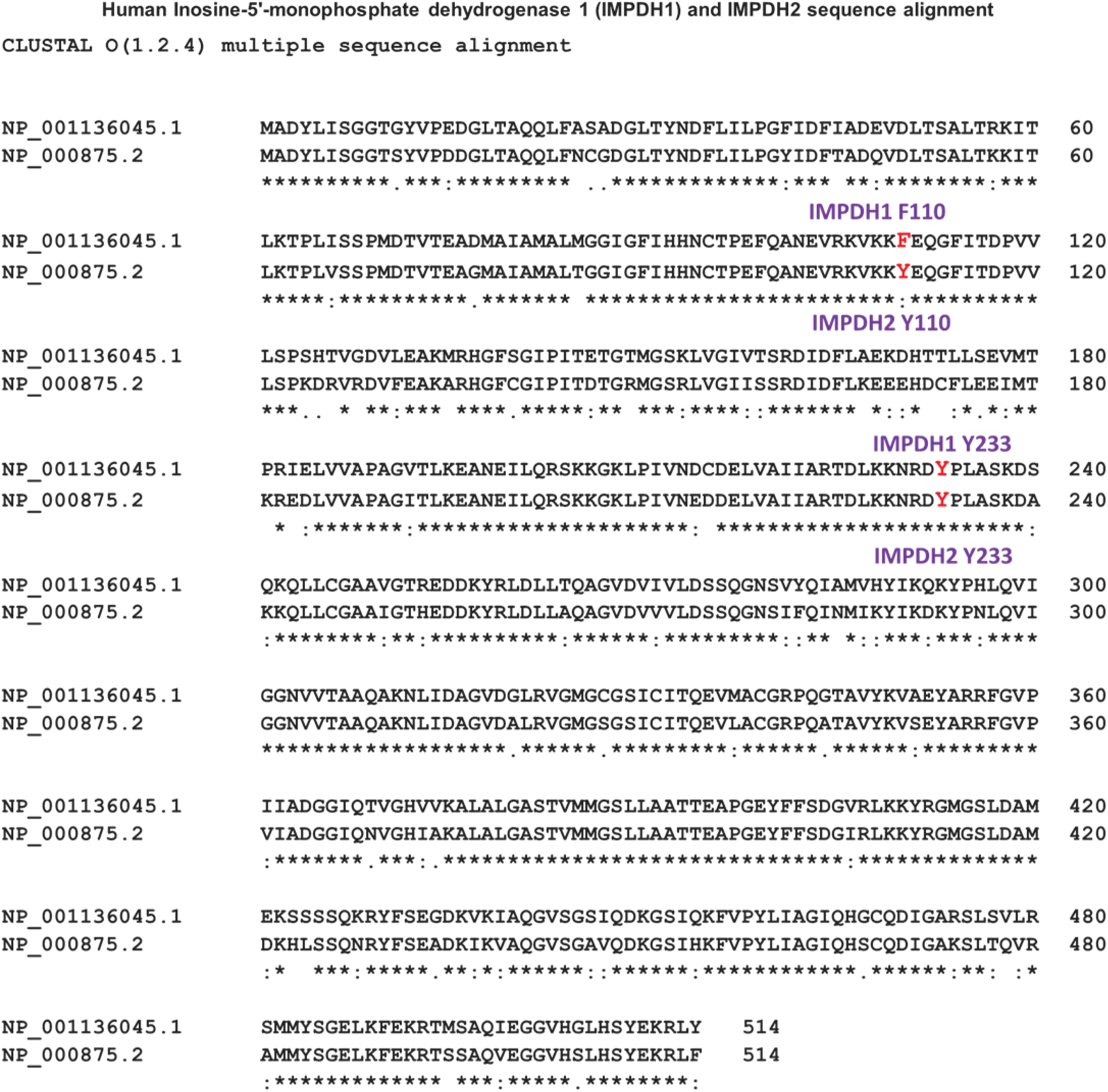
Sequence alignment of human IMPDH1 and IMPDH2.

**Supplementary Fig. 2.**
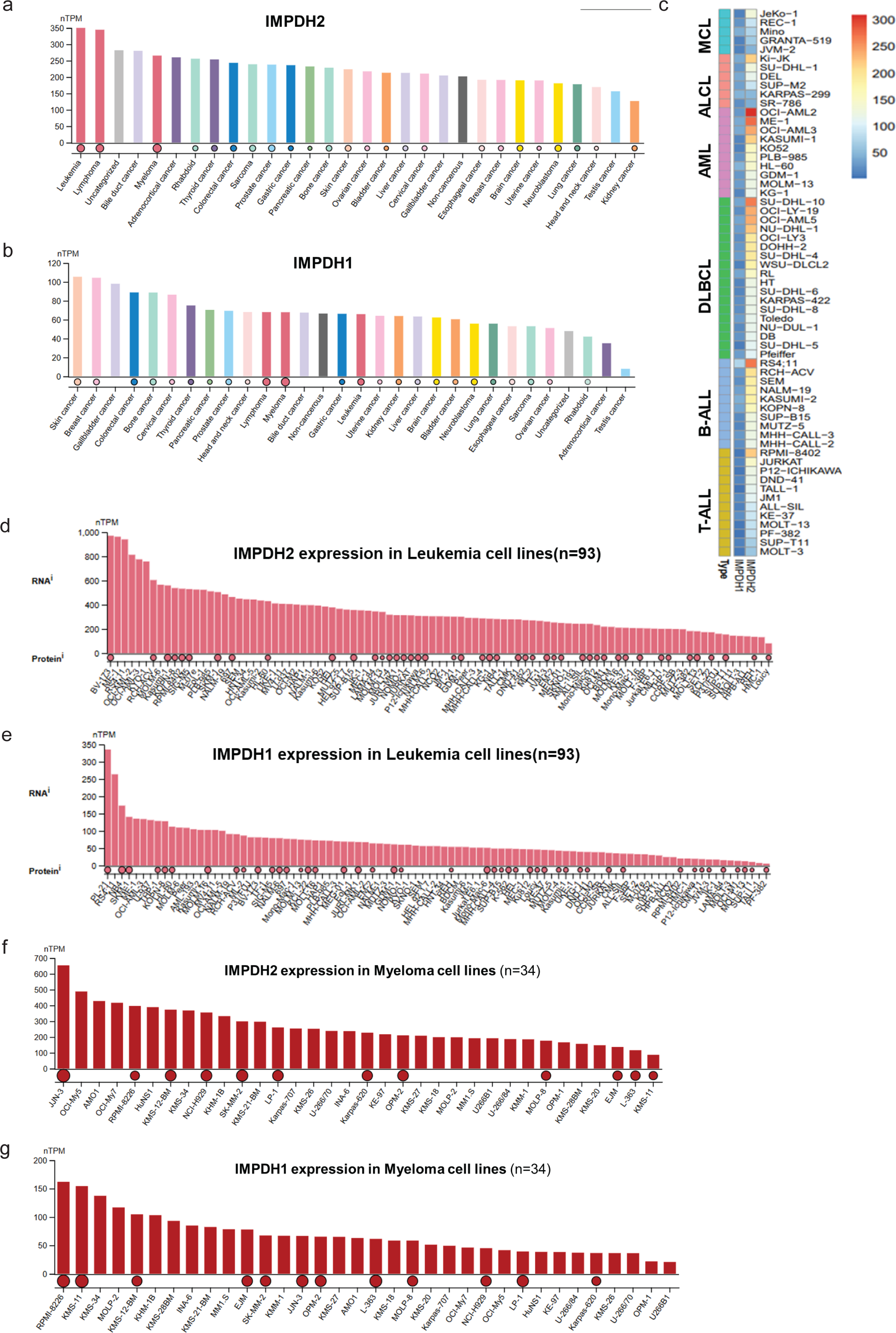
IMPDH2 is broadly overexpressed across human cancers. Transcriptomic analysis of IMPDH2 expression in publicly available datasets reveals significant overexpression in a wide range of malignancies, including both solid tumors and hematological cancers. Elevated IMPDH2 levels are observed in lymphomas, leukemias, colorectal, lung, breast, and ovarian cancers compared to matched normal tissues. These findings underscore the potential role of IMPDH2 in cancer progression and support its candidacy as a therapeutic target across diverse tumor types. **a**, IMPDH2 expressions in various cancers. **b**, IMPDH1 expressions in various cancers. **c**, IMPDH1 and IMPDH2 expressions in various hematological malignancy cell lines. **d, e**, IMPDH2 and IMPDH1 expressions in leukemia cell lines. **f, g**, IMPDH2 and IMPDH1 expressions in myeloma cell lines. This data was curated using the Human Protein Atlas website and TCGA databases.

**Supplementary Fig. 3.**
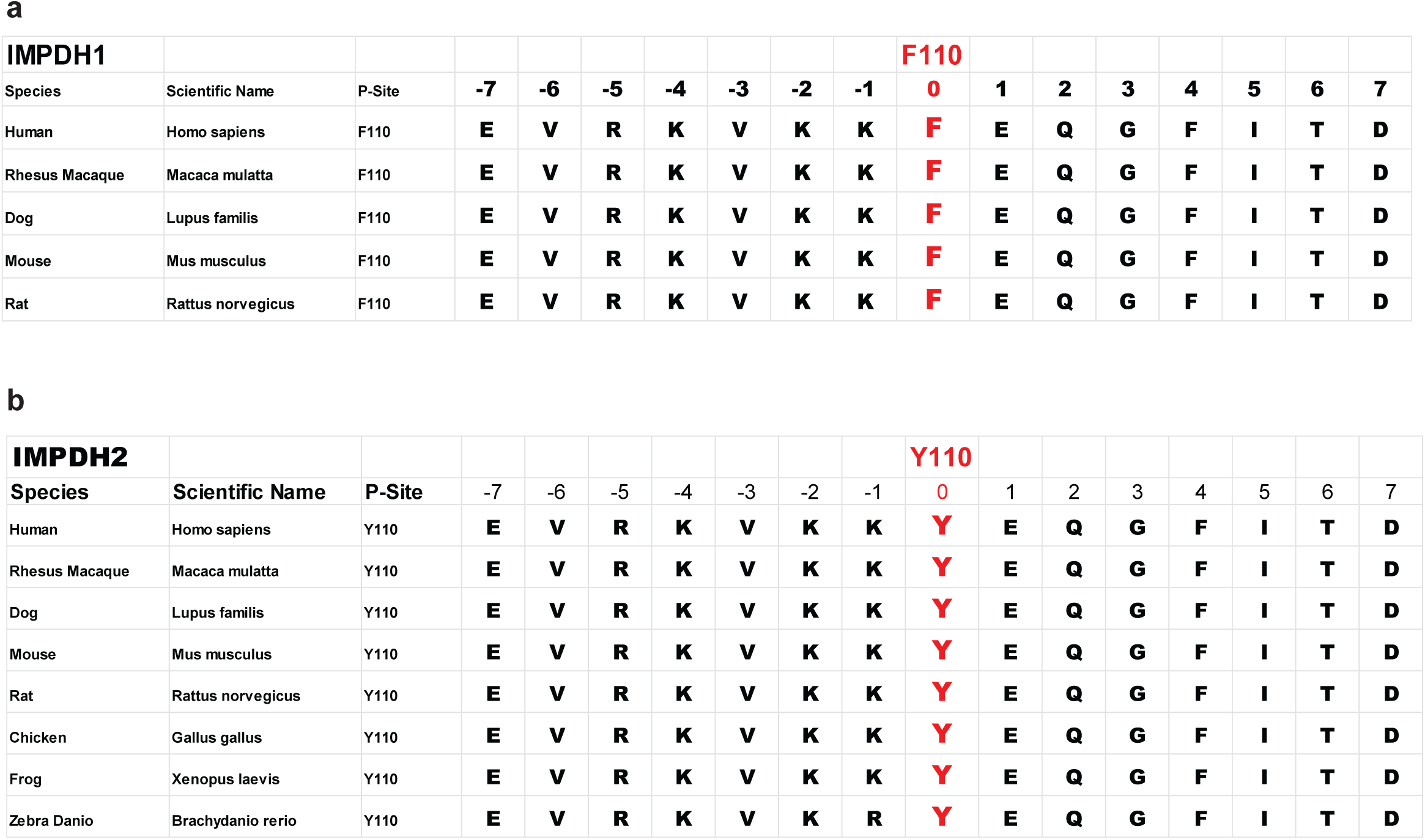
IMPDH1 F110 and IMPDH2 Y110 CBS-1 domain sequence homology of various species.

**Supplementary Fig. 4.**
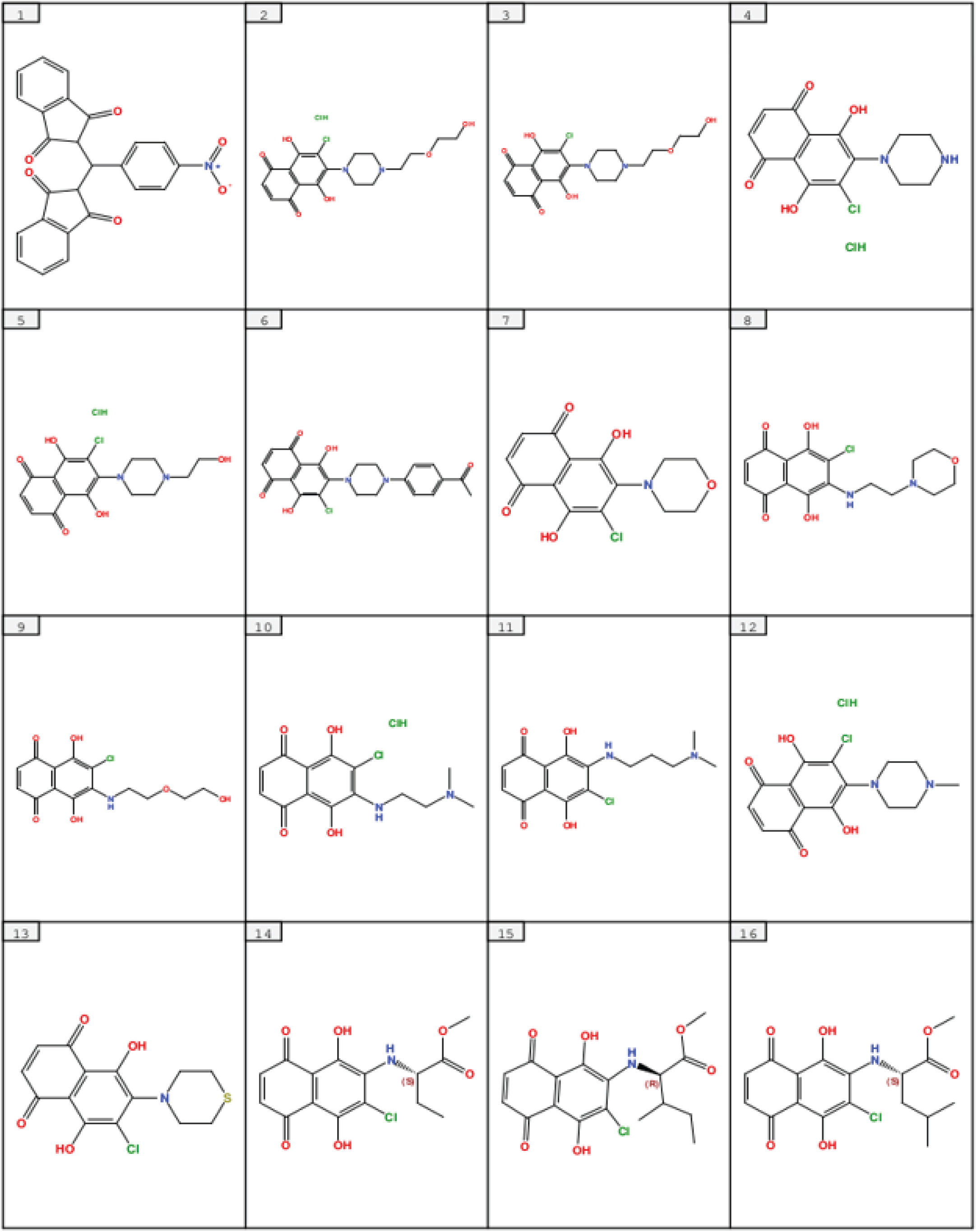
Structure-based in-silico screening and identification of novel IMPDH2 inhibitor analogs.

**Supplementary Fig. 5.**
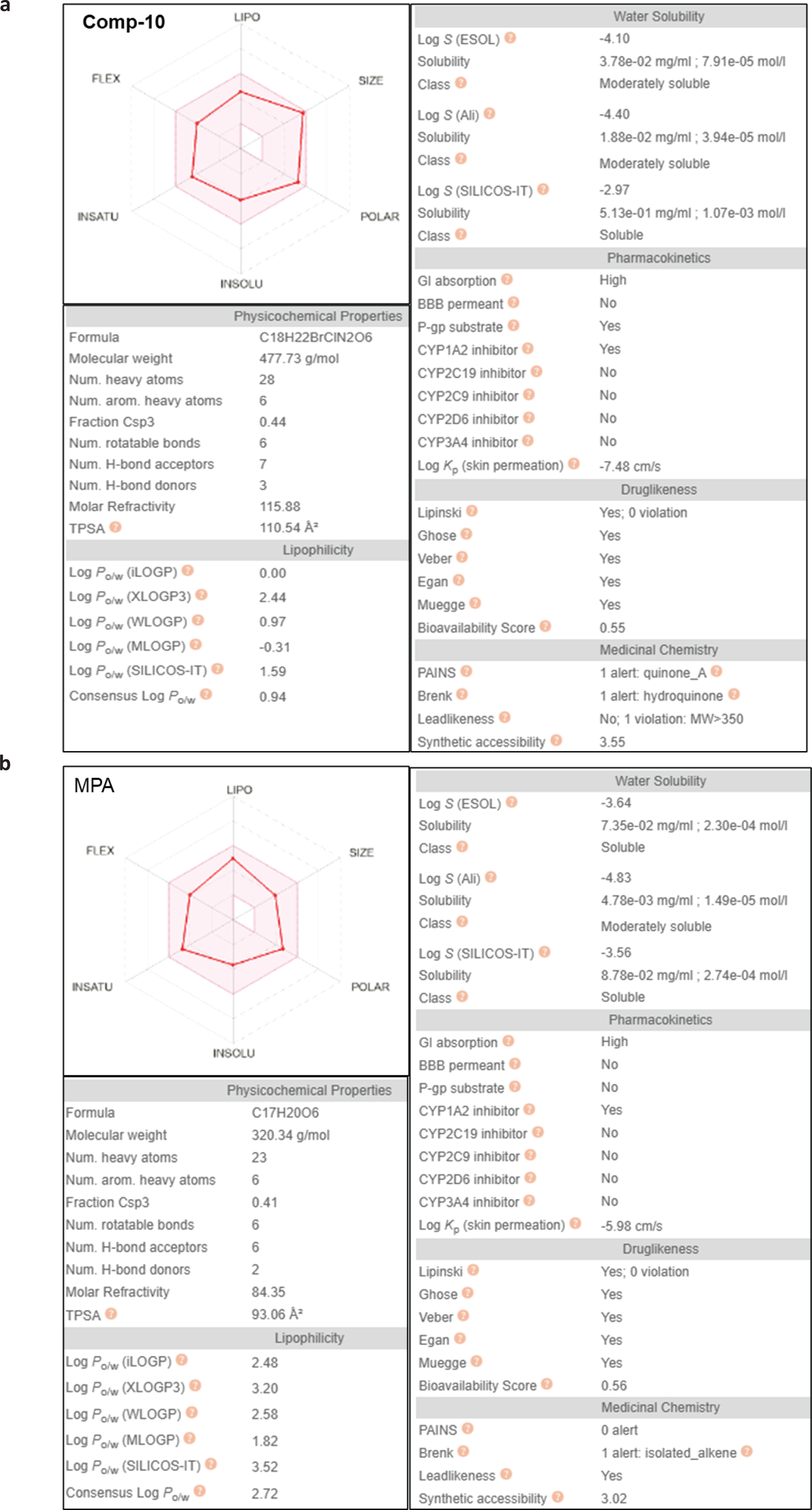
SwissADME analysis predicts favorable drug-likeness and oral bioavailability of Comp-10. **a-b**, Radar plot summarizing six key physicochemical properties of Comp-10 evaluated by SwissADME: lipophilicity (XLOGP3), molecular weight (MW), polarity (topological polar surface area, TPSA), solubility (log S), saturation (fraction of sp³-hybridized carbons), and molecular flexibility (number of rotatable bonds). The pink area denotes the optimal range for oral bioavailability. Comp-10 (MW = 477.73 g/mol, TPSA = 110.54 Å², log S = –4.10, sp³ fraction = 0.44, 6 rotatable bonds) falls within the favorable zone for all six parameters, indicating strong drug-like characteristics and predicted oral bioavailability.

